# 6-Phosphogluconate Dehydrogenase Links Cytosolic Carbohydrate Metabolism to Protein Secretion

**DOI:** 10.1101/480129

**Authors:** Haoxin Li, Maria Ericsson, Bokang Rabasha, Bogdan Budnik, Bridget Wagner, Levi A. Garraway, Stuart L. Schreiber

## Abstract

The proteinaceous extracellular matrix (ECM) is vital for cancer cell survival, proliferation, migration, and differentiation. However, little is known regarding metabolic pathways required in the ECM secretion process. By using an unbiased computational approach, we searched for enzymes whose suppression may lead to disruptions in protein secretion. Here, we show that 6-phosphogluconate dehydrogenase (PGD), a cytosolic enzyme involved in carbohydrate metabolism, is required for endoplasmic reticulum (ER) structural integrity and protein secretion. Chemical inhibition or genetic suppression of its activity led to cell stress accompanied by significantly expanded ER volume and can be rescued by compensating glutathione supplies. Our results also suggest that this characteristic ER-dilation phenotype may be a general marker indicating increased ECM protein congestion inside cells and decreased secretion. Thus, PGD exemplifies a nexus of cytosolic carbohydrate metabolism and protein secretion.

## INTRODUCTION

The ECM, known as the core matrisome, is composed of secreted protein components with diverse chemical or physical properties and undergoes dynamic remodeling by surrounding tissues (Bonnans et al., 2014). Besides structural support, ECM proteins also integrate complex signals to cells with spatial regulation (Hynes, 2009). Copious amounts of ECM proteins are secreted by some cancers including invasive melanoma (Rocco et al., 2011). Their composition, structure, and quantity is central for cancer cell survival, proliferation, and migration (Lu et al., 2012; Pickup et al., 2014).

Given the essentiality of ECM proteins, cells utilize distinct molecular chaperones to guide secretory protein folding (Braakman and Hebert, 2013; Ellgaard and Helenius, 2003). Misfolded proteins that fail quality control are often exported from ER and degraded by the ubiquitin-proteasome machinery (Olzmann et al., 2013). However, it is unclear whether cytosolic metabolic activities might influence ER protein folding and secretion.

In this study, by using a computational screen approach, we identified PGD, a cytosolic carbohydrate-metabolizing enzyme, to be an essential component required for ECM protein secretion. With tool compounds targeting different steps in the protein secretion pathway, we also unveiled ER dilation as a phenotypic indicator of increased secretory protein congestion inside cells and decreased exocytosis.

## RESULTS

### Dependency and Transcription-signature Analysis Reveals Multiple Gene Hits Related to Protein Secretion

To find genes that might be essential for protein secretion, a combination of two complementary computational approaches was applied (Figure 1A, also see STAR Methods). We first searched for candidate genes by correlating their dependency profiles across a panel of cell lines (Wang et al., 2017). Here, we leveraged a genome-wide RNAi screening dataset including 17,098 gene-dependency profiles (DEMETER scores) of 467 screened cell lines (Tsherniak et al., 2017). Given the known central roles of the coatomer (COPI) complex in protein transport mediated by coated vesicles (Cosson and Letourneurt, 1997), we hypothesized genes that correlated with dependency profiles of the coatomer complex might also be relevant in protein secretion. We chose the *COPA* dependency vector as a representative, given different coatomer components (e.g., *COPA*, *COPB1*, *COPB2*, etc.) have similar profiles (Figure S1A). As an output, candidate genes were ranked by their Pearson correlations with the *COPA* dependency vector.

**Figure 1.**
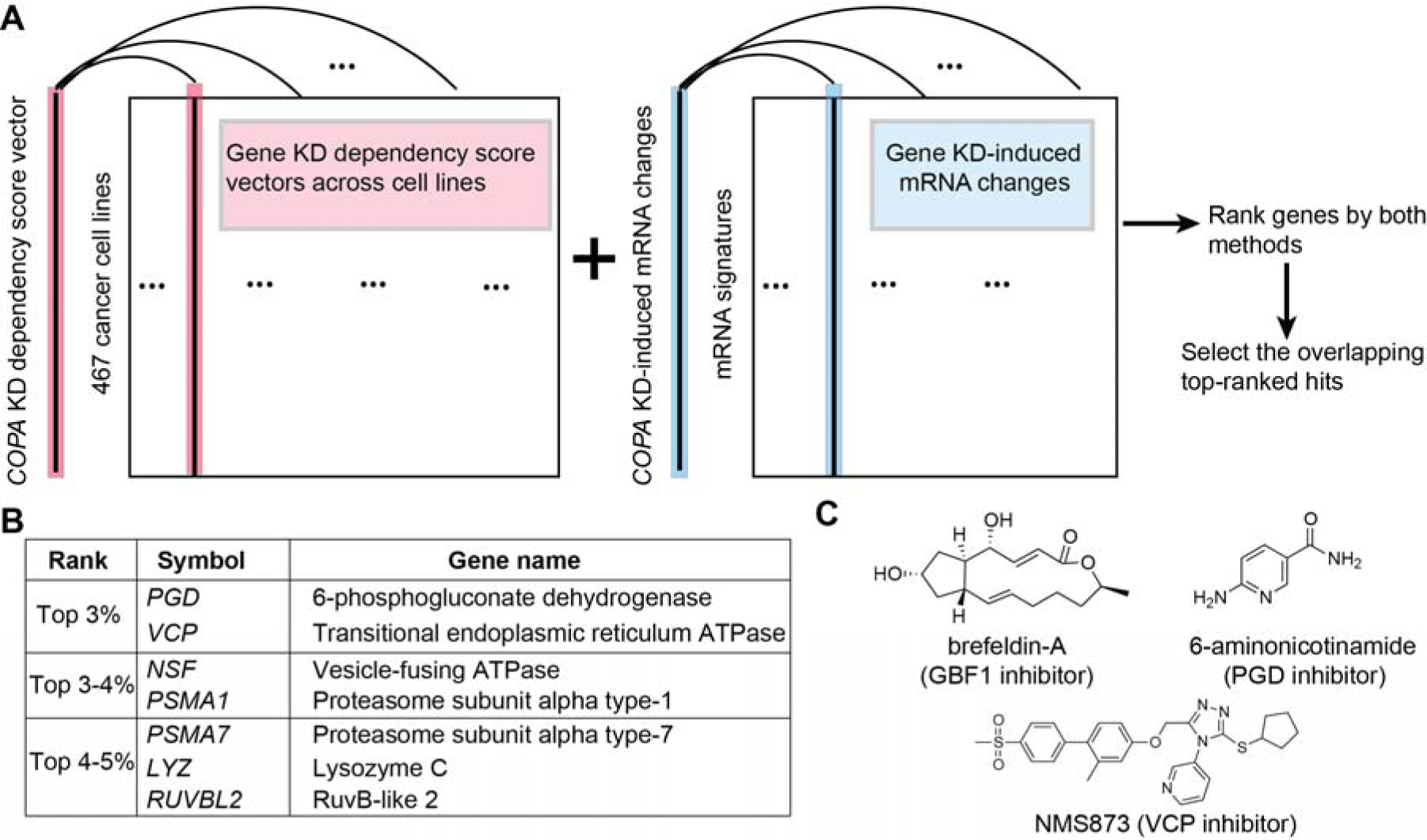
Dependency and Transcription-signature Analysis Reveals Multiple Gene Hits Related to Protein Secretion. (A) A combination of two computational approaches was used to discover potential genes essential for protein secretion. The *COPA* dependency vector was correlated with other dependency vectors (shown as columns) in a genome-wide RNAi screen dataset including a panel of 467 cancer cell lines. The *COPA* KD-induced mRNA signature was compared with other available gene KD-induced mRNA signatures (shown as columns). The top ranked genes in both methods were then selected as hits. (B) Top-ranked enzyme-coding gene hits filtered by analysis in (A). The threshold was set to be 3%, 4%, 5% and the hits were ranked by order. (C) Structures and mechanisms of compounds used in this study. See also Figure S1.

Next, we looked for perturbations eliciting similar gene expression alterations in response to secretory pathway blockade and hypothesized that such connected perturbations might confer related physiological effects on cells (Lamb et al., 2006). Currently, there are 3799 gene knock-down (KD)-induced transcriptional signatures in the Connectivity Map database (Subramanian et al., 2017). As an output, candidate genes were ranked according to their similarity with *COPA* in terms of KD-induced transcriptional changes.

To combine results from both computational analyses, we filtered the top hits with cutoff at different percentiles and retained those genes that were overlapping (Table S1). Here, we focused on hits having enzymatic activities, which include 6-phosphogluconate dehydrogenase (PGD) and transitional endoplasmic reticulum ATPase (VCP) (Figure 1B). In the oxidative phase of pentose phosphate pathway, NADP-dependent PGD catalyzes the decarboxylation of 6-phosphogluconate (6PG), a metabolite derived from glucose oxidation, yielding ribulose 5-phosphate and NADPH (Figure S1B). PGD is inhibited by the tool compound 6-aminonicotinamide (6AN) (Figure 1C), which is metabolized to an analog of NADP (Köhler et al., 1970). VCP is an ATPase located on the cytosolic face of the endoplasmic reticulum (ER), which is essential for exporting misfolded proteins destined for proteasome degradation (Bodnar and Rapoport, 2017) and can be inhibited by a potent allosteric inhibitor NMS873 (Magnaghi et al., 2013) (Figure 1C). Taken together, our computational analysis revealed *PGD* and *VCP* as candidate genes related to protein secretion.

### PGD Inhibition Causes Significant Decreases of Protein Secretion to the Extracellular Space Accompanied by ER dilation

To test the effects of PGD and VCP inhibition in cells, we chose WM793 as a sensitive melanoma model cell line (Figure S2A). As a positive control, brefeldin A was included to block protein secretion by inhibiting guanine nucleotide exchange factor (GBF1) and disrupting coatomer coat assembly (Helms and Rothman, 1992; Misumi et al., 1986). Interestingly, we found accumulation of vacuole-like structures in cells treated with brefeldin A, 6AN, or NMS873 (Figure 2A). To investigate the fine structural details of these cytoplasmic vacuoles, we examined untreated and treated cells using transmission electron microscopy (TEM). We observed flattened, membrane-enclosed ER sacs in control cells; however, a significant proportion of those treated by brefeldin A, 6AN, or NMS873 showed ER dilation (Figure 2B). This phenotype was absent in staurosporine-induced apoptosis or rapamycin-induced autophagy (Figure S2B).

**Figure 2.**
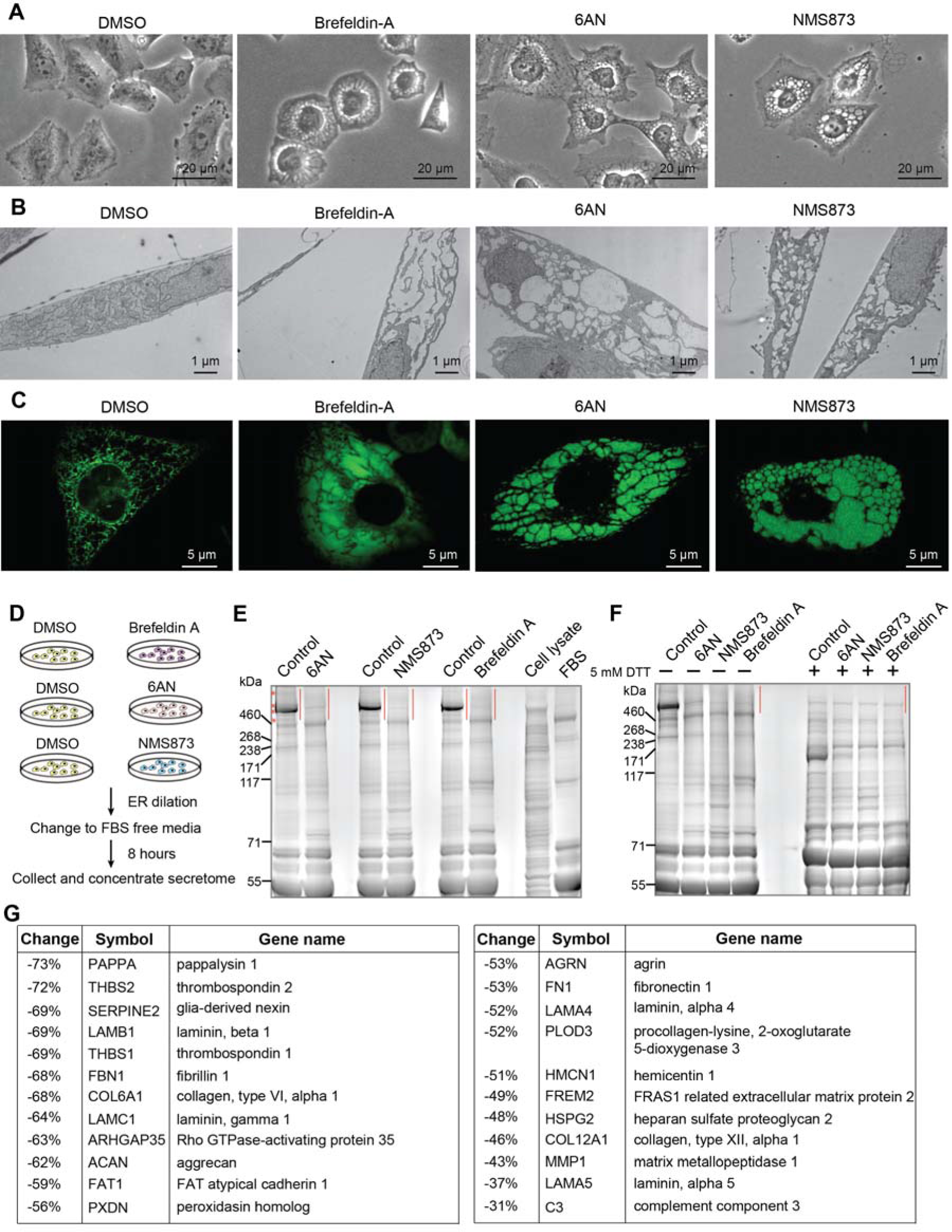
PGD Inhibition Causes Significant Decreases of Protein Secretion to the Extracellular Space Accompanied by ER dilation. (A-B) Representative WM793 cells under (A) light microscope and (B) electron microscope. The treatments include brefeldin A (1 μM, 36 h), 6AN (20 μM, 48 h), and NMS873 (5 μM, 18 h). (C) Representative immune-fluorescence images of GFP-tagged calreticulin in live U2OS cells. The treatments include brefeldin A (1 μM, 36 h), 6AN (20 μM, 72 h), and NMS873 (5 μM, 18 h). (D) Secretome analysis workflow of WM793 cells with/without induced ER dilation. (E) A representative gel image showing secreted proteins separated by SDS-PAGE (non-reducing condition) and stained with Coomassie blue. Untreated whole cell lysates and FBS were included as additional controls. Several bands present in DMSO-treated controls but absent under treatments causing ER dilation are marked with asterisks. Proteins in gel bands highlighted in red were subsequently digested and peptides were labeled with tandem mass tags for quantitation. (F) Same as (E) except a reducing condition with 5 mM DTT was tested. (G) Average secreted protein abundance changes with ER dilation induced by brefeldin A, 6AN, or NMS873 versus DMSO-treated control. See also Figure S2.

Next, we asked whether *PGD* loss leads to ER dilation. CRISPR-Cas9 was used to generate isogenic WM793 cell lines with inducible *PGD* knockout (KO), and RNAi was used to create cell lines with stable *PGD* knockdown (KD). In both systems, reductions in PGD protein abundance were observed (Figure S2C) and among these cells, a significant proportion displayed markedly dilated ER (Figures S2D and S2E) and reduced proliferation (Figures S2F and S2G).

To confirm the organelle origin and to monitor ER structural alteration in live cells, we used a U2OS cell line genetically engineered to express GFP-calreticulin, an ER lumen marker, at endogenous levels (Wang et al., 2016). By using confocal microscopy, we observed that ER lumen was markedly dilated in cells treated with 6AN, NMS873, or brefeldin A (Figure 2C).

To investigate the relationship between ER dilation and protein secretion, we collected and analyzed secreted proteins from cells with ER dilation or control treatment (Figure 2D). Briefly, when cells started to have ER dilation (using 6AN, NMS873 or brefeldin A), the culture media was refreshed without fetal bovine serum (FBS). After 8 hours of incubation, proteins in the conditioned media were concentrated and separated by SDS-PAGE (Figure 2E). Considering potential contamination by residual FBS or intracellular proteins released following cell death, we also included FBS and whole cell lysates as additional controls. Our results showed that cells with ER dilation induced by 6AN, NMS873, or brefeldin A had significantly decreased protein secretion in the 460-600 kDa range under non-reducing conditions. By reducing samples with dithiothreitol (DTT), these bands vanished (Figure 2F), suggesting the major protein contents were rich in disulfide bonds.

We next processed the proteins in the range of 460-600 kDa and quantified the digested peptides with LC-MS to identify the affected secreted proteins. ER dilation induced by the three different compounds (6AN, NMS873, and brefeldin A) led to surprisingly similar reductions in various secretory proteins, including thrombospondin (THBS1, THBS2), lamin (LAMB1, LAMC1, LAMA4), fibrillin 1 (FBN1), collagen (COL6A1, COL12A1), fibronectin (FN1), secreted enzymes (PAPPA, PLOD3, MMP1), and other ECM proteins (Figure 2G). Taken together, these results suggest that PGD or VCP suppression leads to decreased protein secretion into the extracellular space and is accompanied by a cell-stress phenotype characterized by ER dilation.

### ER Dilation is Associated with Accumulation of Secretory Proteins Inside Cells

We considered two hypotheses that explain the observed reduction in protein secretion. First, cells under stress might have reduced synthesis of secretory proteins. Alternatively, the synthesized secretory proteins might accumulate inside cells without being exported. To test which is operable, we examined the cellular proteomic changes induced by 6AN or NMS873 during ER dilation in WM793 cells (Figure 3A). We were able to quantify 5629 proteins in the 6AN group and 6029 proteins in the NMS873 group (Table S2). We asked how similar were the proteome-wide abundance changes in either group. Remarkably, the proteins changed in response to 6AN treatment were significantly correlated with those changed in response to NMS873 (Pearson correlation: 0.61, p<10-10) (Figure 3B), indicating similar influences of these two treatments on a proteome-scale and echoing the initial computational analysis of signature similarity. To obtain unbiased classification of the significantly changed proteins based on their subcellular localization, we applied Gene Set Enrichment Analysis (GSEA) (Subramanian et al., 2005) to the individual proteins ranked by their fold changes and ordered the results (GO cellular component ontology) by normalized enrichment scores (Figures 3C and 3D). The significantly enriched proteins in both 6AN- and NMS873-treated cells mainly came from ER lumen, transport vesicle, and extracellular matrix, all of which are topologically equivalent in the secretory pathway (Figure 3E and Table S2). This evidence strongly supports the hypothesis that secretory proteins are produced but trapped inside cells instead of being released during ER dilation.

**Figure 3.**
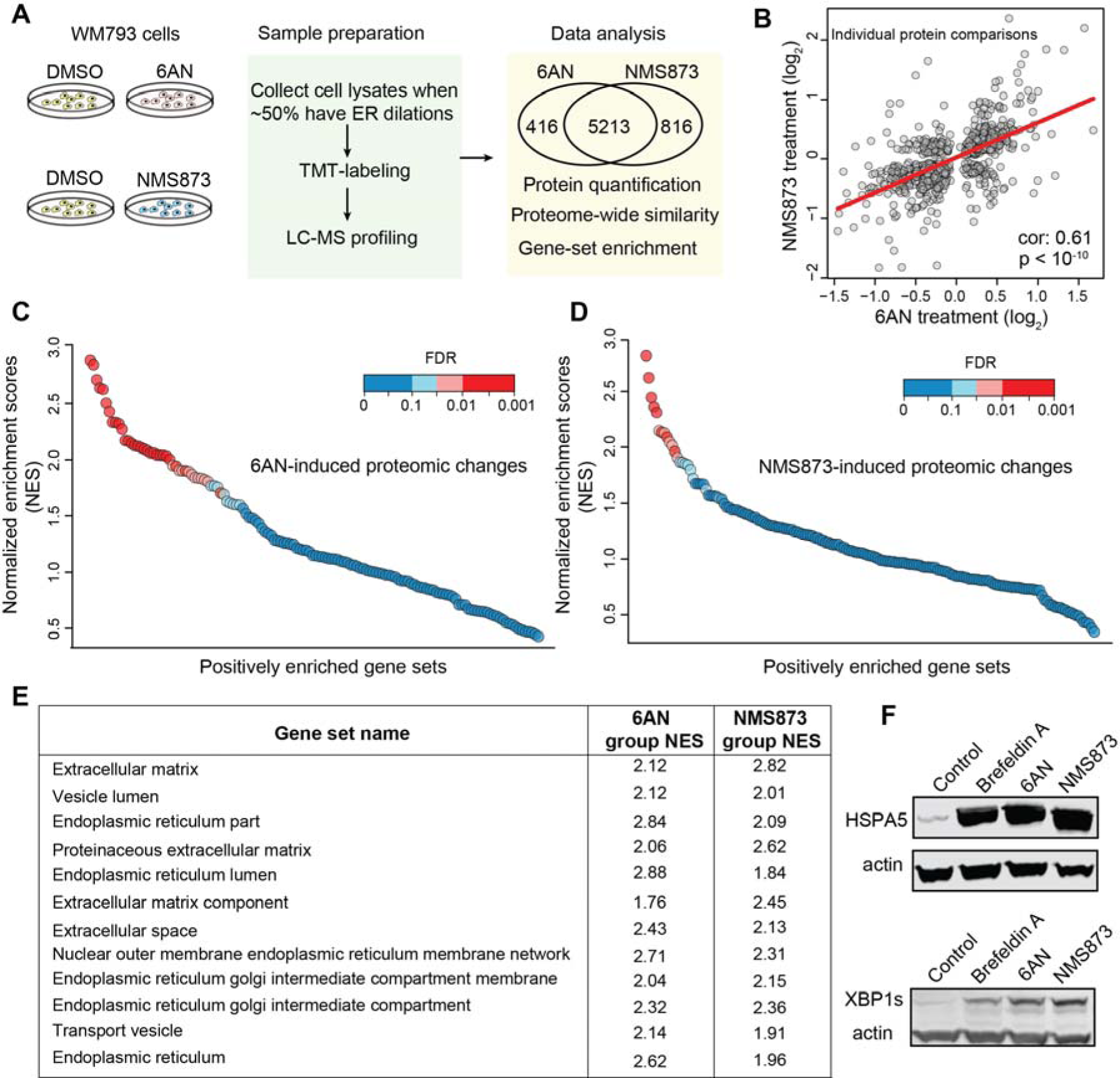
ER Dilation is Associated with Accumulation of Secretory Proteins Inside Cells. (A) Proteomic analysis workflow of WM793 cells with induced ER dilation or control. (B) Comparisons of 6AN-induced protein abundance changes versus control and NMS873-induced changes versus control. Each dot represents an up-regulated or down-regulated protein (p<0.05). The Pearson correlation was 0.61 with p<10^-10^. (C) List of positively enriched gene sets (GO cellular component ontology) based on 6AN-induced proteomic changes. (D) List of positively enriched gene sets (GO cellular component ontology) based on NMS873-induced proteomic changes. (E) Enriched gene sets that are significant (FDR<0.05) in both (C) and (D). (F) Immunoblots of HSPA5 (Bip) and spliced XBP1 (XBP1s) in WM793 cells with different treatments including brefeldin A (1 μM, 36h), 6AN (20 μM, 48h), and NMS873 (5 μM, 18h). See also Figure S3.

Generally, only properly folded proteins can exit ER and proceed to secretion (Ellgaard and Helenius, 2003). To test the overall protein folding status in the ER we evaluated expression of HSPA5, an ER-located chaperone for misfolded proteins, and the spliced form of XBP1, a key transcriptional regulator controlling cytoprotective responses, and found both to be strongly induced in cells with ER dilation (Figures 3F and S3A). Furthermore, we observed that both PGD and VCP suppression induced an up-regulation of UGGT1 (Figure S3B), which senses and reglucosylates ER proteins with defects to continue the calnexin/calreticulin chaperone cycle (Ritter and Helenius, 2000). We also found increased levels of regulators involved in recognizing, ubiquitinating, and exporting misfolded secretory proteins for degradation – including HYOU1 (Steel et al., 2004), DNAJB9 (Dong et al., 2008), ERLEC1 (also known as XTP3-B or erlectin) (Christianson et al., 2008), SEL1L, UBE2J1, and SYVN1 (also known as HRD1 or synoviolin) (Burr et al., 2010) (Figures S3C-H). Together, these results suggest that increased misfolded proteins accumulate inside the dilating ER to a degree that triggered unfolded protein responses (UPR) including up-regulation of cellular machineries facilitating protein folding or clearance.

### PGD Suppression Induces ER Dilation by Decreasing GSH Supplies

Next, we asked what metabolic feature correlates with cell-line sensitivity to PGD suppression. We leveraged the comprehensive Cancer Cell Line Encyclopedia (CCLE) metabolomic database (Li et al., 2018) and correlated 225 metabolite features with the PGD dependency profile in the cell line panel (Figure 4A). We found that those cell lines with higher baseline levels of 6-phosphogluconate (6PG) also had greater dependency on PGD (p < 10^-10^) (Figure 4B), suggesting the use of 6PG to identify cancer cells that are sensitive to PGD suppression.

**Figure 4.**
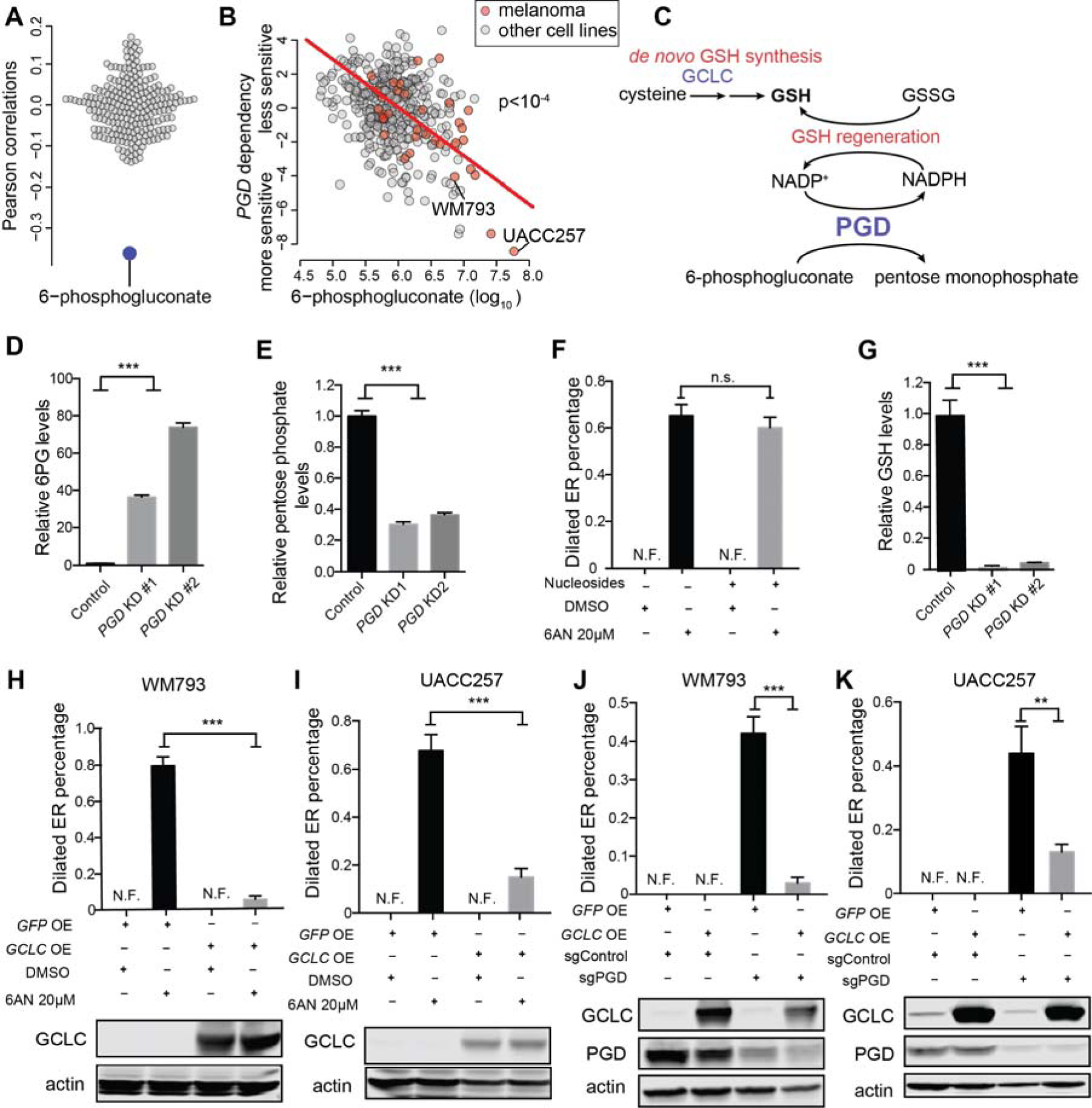
PGD Suppression Induces ER Dilation by Decreasing GSH Supplies. (A) Among 225 metabolites measured, 6-phosphogluconate is the most associated metabolite with *PGD* dependency across 417 overlapping cancer cell lines. Each dot represents a metabolite. (B) Basal 6-phosphogluconate accumulation correlates with greater sensitivity to PGD KD in different cancer cell lines (represented as points). (C) Schematic depicting PGD metabolism and related metabolic pathways maintaining reduced glutathione (GSH) levels. (D) Relative 6-phosphogluconate levels in WM793 cells with *PGD* KD versus control. (E) Relative pentose monophosphate levels in WM793 cells with *PGD* KD versus control. (F) Effects of nucleoside supplements (30 μM cytidine, guanosine, uridine, adenosine, and thymidine) on 6AN-indudced ER dilation in WM793 cells (48 h). (G) Relative GSH levels in WM793 cells with *PGD* KD versus control. (H-I) Effects of *GCLC* overexpression on the percentages of cells with ER dilation induced by 6AN in (H) WM793 cells, 2 days of treatment and (I) UACC257 cells, 3 days of treatment. N.F., dilated ER not found. (J-K) Effects of *GCLC* overexpression on the percentage of cells with ER dilation upon *PGD* KO in (J) WM793 cells and (K) UACC257 cells, 9 days after 1 μg/ml doxycycline treatment. The bar plots represent mean ± SEM (*** p < 0.005; ** p < 0.01; * p < 0.05; n.s., non-significant, p > 0.05; n = 4) (D-K) See also Figure S4.

To explain how PGD suppression induces a strong ER dilation phenotype, we probed the metabolomic changes using LC-MS. We first showed that PGD knockdown induced about 50-fold increase of its substrate, 6-phosphogluconate (Figures 4C and 4D), suggesting lowered enzymatic activity. Additionally, the pentose phosphate metabolites synthesized by PGD and the downstream nucleoside triphosphate species were reduced more than half by PGD knockdown (Figures 4E and S4A). We therefore hypothesized that if reduced nucleoside supplies cause ER dilation, supplementing them in culture may reduce ER dilation. However, no noticeable rescue effect was observed with chemical or genetic suppression of PGD (Figures 4F and S4B).

PGD generates NADPH in the decarboxylation reaction, a reducing equivalent that is essential for redox homeostasis including glutathione regeneration. Consistently, we observed depleted reduced glutathione (GSH) in cells with PGD KD (Figure 4G). We therefore hypothesized that if this GSH depletion caused ER dilation, increased expression of glutamate-cysteine ligase (GCLC), an enzyme that controls the rate-limiting step of glutathione *de novo* synthesis (Figure 4C), would suppress PGD inhibition-induced ER dilation. Accordingly, GCLC overexpression effectively reduced the percentage of cells with dilated ER after 6AN treatment (Figures 4H, 4I, and S4C) or doxycycline-induced PGD knockout (Figures 4J and 4K) in WM793 and UACC257 cells.

Earlier studies reported that a decrease in GSH availability led to lipid peroxidation and consequent ferroptosis (Dixon et al., 2012). To test whether lipid peroxides caused ER dilation, we evaluated a small-molecule tool compound ferrostatin that reduces lipid peroxidation. We found that ferrostatin rescued lipid peroxidation-induced ferroptotic cell death (Figure S4D), but did not reduce ER dilation caused by PGD suppression (Figure S4E). Collectively, these results suggest that PGD suppression induces ER dilation by limiting GSH availability in a lipid peroxidation-independent pathway.

## DISCUSSION

Dysregulation of ECM protein exocytosis has been implicated in various human pathologies and diseases including cancer (Aridor and Hannan, 2000; Bonnans et al., 2014; Lin et al., 2008). The survival, proliferation, and metastasis of cancer cells require a tailored ECM niche (Lu et al., 2012; Pickup et al., 2014). Previous studies revealed various modules involved in ER protein sorting and quality control (Ellgaard and Helenius, 2003). However, the metabolic environment that supports protein secretion is yet to be characterized. By using an unbiased computational approach, we discovered PGD as an intercompartmental link between cytosolic carbohydrate metabolism and protein secretion. Its chemical or genetic suppression led to an intriguing ER dilation phenotype, similar to that induced by small molecules disrupting coatomer assembly or blocking misfolded protein removal from ER. This PGD-suppression-induced ER dilation is a result of reduced downstream GSH regeneration, which can be rescued by boosting *de novo* GSH synthesis. Using quantitative proteomic profiling, we showed that cells with ER dilation displayed markedly reduced protein secretion into the extracellular space and increased misfolded protein congestion in the secretory pathway. Therefore, our results demonstrate that ECM protein blockade can be triggered by aberrant metabolic activities and provide clues for future studies investigating their causal roles in various disease contexts involving secretion dysregulation.

The inclusion of the small-molecule probes brefeldin A and NMS873 based on our initial computational analysis provided clues to explain the ER dilation phenotype induced by PGD suppression. As a positive control, brefeldin A is a well-validated tool compound used to block coat assembly on secretory vesicles and therefore inhibit protein secretion (Helms and Rothman, 1992; Misumi et al., 1986). NMS873 inhibits VCP, an ATPase that removes misfolded ER proteins destined for cytosolic proteasome degradation through a channel (Bodnar and Rapoport, 2017; Ye et al., 2001). These two compounds blocked either normal secretory protein exit or misfolded proteins clearance. We therefore speculate that when these secretory products overwhelm the ER protein folding/clearance capacity, lumen pressure increases and in turn causes volume expansion.

Similarly, PGD suppression, which decreases protein secretion and increases protein retention, may also induce ER dilation by creating protein-cargo congestion. Given that ER is specialized for disulfide bond formation and rearrangements with carefully maintained redox balance (Chakravarthi et al., 2006), PGD-suppression-induced GSH imbalance might cause non-native disulfide bond formation and thus result in accumulation of incorrectly folded proteins in the lumen (Chakravarthi and Bulleid, 2004; Molteni et al., 2004). Unlike VCP inhibition, we speculate that the PGD-induced congestion is driven by increases in upstream protein misfolding instead of decreases in downstream clearance. Although it is currently unknown how the cytosolic GSH is imported into the ER, our data underscore the importance of PGD in supporting secretory protein folding by maintaining GSH balance and suggest a “congestion-to-dilation” model.

## SIGNIFICANCE

Our work illustrates that PGD, an enzyme in the pentose phosphate pathway links cytosolic carbohydrate metabolism to protein secretion, highlighting the importance of cytosolic metabolic activities in supporting ER functions. We also demonstrate an unbiased computational approach to unveil distinct genes involved in inherently related biological processes. Perturbations of these nodes using small-molecule tool compounds have analogous phenotypic impact. This dependency and transcription-signature analysis takes advantage of in-depth molecular characterization of cancer cell lines and is a powerful way to discover perturbations that induce illuminating phenotypes.

## Supporting information

## ACKNOWLEDGEMENT

The authors would like to thank Christina Woo and Paul Schwein for valuable suggestions related to proteomic analysis. We thank Tom Rapoport, Yilong Zou, Craig Strathdee, Marios Giannakis, Shawn Nelson, and William Sellers for helpful discussions. We acknowledge Jeffrey Wyckoff from the Koch Institute for help with confocal experiments. This work is supported by the NCI’s Cancer Target Discovery and Development (CTD2) Network (grant number U01CA217848, awarded to S.L.S.) and CCLE grant from Novartis. Haoxin Li is a fellow in the Herchel Smith Graduate Fellowship Program.

## AUTHOR CONTRIBUTIONS

H.L. designed the experiments, performed research, and was supervised by L.A.G and

S.L.S. H.L. and M.E. did the TEM experiments. B.R., C.S., and B.B. assisted with other experiments. H.L. and S.L.S. wrote the manuscript in discussion with other co-authors.

## TABLE WITH TITLES and LEGENDS

Table S1. Gene Hits Revealed by Dependency and Transcription-signature Analysis, Related to Figure 1

Table S2. Whole Proteome Profiling of WM793 Cells with Induced ER Dilation, Related to Figure 3

## STAR METHODS

### CONTACT FOR REAGENT AND RESOURCE SHARING

Further information and requests for resources and reagents should be directed to and will be fulfilled by the Lead Contact, Stuart Schreiber.

### EXPERIMENTAL MODEL AND SUBJECT DETAILS

WM793 (male) and UACC257 (female) cells were maintained in RPMI media (37°C, 5% CO_2_) with 10% (v/v) heat-inactivated fetal bovine serum (FBS) and 0.2% (v/v) antibiotic normocin (InvivoGen). U2OS (female) cells were maintained in DMEM media (37°C, 5% CO_2_) with 10% (v/v) heat-inactivated FBS and 0.2% (v/v) antibiotic normocin. The cell lines were confirmed by SNP genotyping.

### METHOD DETAILS

#### Dependency and Transcription Signature Analysis

This computational analysis aimed to find genes that might be essential for protein secretion and was separated into two parts including dependency signature analysis and transcription signature analysis. First, in a gene dependency dataset based on RNAi screens among 467 cancer cell lines (Tsherniak et al., 2017), each column was termed as a gene dependency vector (the quantitative dependency of that gene among these cell lines). We correlated all 17,098 columns in this dataset with the *COPA* column (representative of the coatomer complex) and ranked the hits based on their Pearson correlations (see Table S1). Second, in the connectivity map database (https://clue.io/cmap), there were 3799 gene knockdown-induced mRNA changes. Such transcription signatures were compared with the *COPA* knockdown-induced signature and the hits were scored as described previously (Subramanian et al., 2017) (see Table S1). After completing these two steps, we next combined the results and searched for overlapping top hits with cutoff at different percentiles. For example, the top 3% hits in both the dependency and transcription analysis results included *VCP* and *PGD*. In this study, we focused on enzyme hits selected by the enzyme commission number annotations from https://www.genenames.org/. The hits were included in Table S1.

#### ER Dilation Percentage Quantification

The percentages of cells with significantly dilated ER under different experimental conditions were visualized and counted manually under a light microscope using the EVOS FL cell imaging system. At least 3 independent fields in each of the biological replicates were used for data acquisition. Representative images were processed following community standards and correctly represented the original data.

#### Metabolite Extraction and LC-MS Analysis

Five days after resistance marker selection (2 μg/ml puromycin), WM793 cells were seeded in 6 well plates at a density of 4*10^5^/well. 2 days later, cells were collected after removal of media and were washed with ice-cold 0.9% NaCl (1.5 ml/well). Plates were then moved to dry ice and 1 ml/well of 80% methanol solution containing 10 ng/ml valine-d8 (internal standard) was added to each sample well. Cell lysates were scraped and transferred to Eppie tubes and were vortexed for 10 min in a cold room. After centrifugation at 10,000 g for 10 min (4°C), the supernatants were dried using Speedvac. The LC/MS-based profiling was performed as described previously (Birsoy et al., 2015).

#### Constructs and Clones

sgRNA for *PGD* KO (guide: CAGGAGGGAACAAAGAAGCG) or *GFP* KO (control) was cloned into pLV709 doxycycline-inducible sgRNA expression vectors (puromycin resistance). Tetracycline-free FBS was used to maintain cells before doxycycline-inducible gene knockout experiments.

For shRNA-mediated knockdown experiments, the shRNAs targeting *PGD* (#1: TRCN0000274974, #2: TRCN0000274976) or *GFP* (control) were cloned into constitutive shRNA expression vectors pLKO-TRC005 (puromycin resistance) by the Broad Institute Genome Perturbation Platform. The *GCLC* cDNA in pDONR221 vector was obtained from Harvard Plasmid ID database (HsCD00296342) and was cloned into pLV406 vector for constitutive expression with standard gateway cloning (G418-resistance).

#### Generation of Isogenic Cell Lines

Vectors expressing sgRNA along with lentiviral packaging and envelope vectors (pCMV-dR8.91 and VSV-G) were transfected into HEK293T cells using the FuGENE 6 transfection reagent (Promega) according to the manufacturer’s protocol. The media were changed to DMEM with 30% FBS 18 hours after transfection. The virus-containing supernatant was collected 48 hours later and passed through a 0.45 μm filter to eliminate cells. Target cells were seeded in 6-well tissue culture plate at a density about 2*10^5^/well and were infected in media containing 8 μg/ml of polybrene. Spin infection was performed by centrifugation at 2,200 rpm for 1 hour. One day after infection, the virus-containing media was removed and media containing selection antibiotic was added (enough to kill cells without resistance markers). The gene suppression or expression efficiencies were validated using immunoblotting.

#### Immunoblotting

Cells were trypsinized, pelleted, and then washed twice in cold PBS. They were lysed in RIPA buffer on ice for 20 min. Incompletely lysed cells and debris were precipitated at 16,000 g for 20 min in a microcentrifuge at 4 °C. The protein concentration was estimated with bicinchoninic acid (BCA) assay. Equal amounts of samples were mixed with loading buffer (Bio-rad) and were resolved by 4-12% Bis-Tris SDS-polyacrylamide gel electrophoresis (SDS-PAGE). Proteins were then transferred to PVDF membranes using the iBlot system (Invitrogen). The membrane was blocked for 1 hr in Li-Cor Odyssey blocking buffer (LI-COR) and incubated in primary antibodies overnight at 4 °C. Following 3 times of 10 min washes in Tris-buffered saline (pH 7.4) with 0.1% Tween-20 (TBST), the membrane was incubated with secondary antibodies for 1 hour. It was then washed again in TBST for three times (10 min each) with protection from light and scanned using the Odyssey imaging system (LI-COR). The antibodies used include: anti-PGD (Santa Cruz, sc-398977, 1:100), anti-GCLC (Abcam, ab190685, 1:5000), anti-actin (Cell Signaling Technology, #3700, #8457, 1:2000), anti-HSPA5 (Cell Signaling Technology, #3177, 1:1000), anti-XBP1s (Cell Signaling Technology, #12782, 1:1000), IRDye® 800CW goat anti-mouse IgG (H+L) (LI-COR, 925-32210, 1:10000), and IRDye® 680RD goat anti-rabbit IgG (H+L) (LI-COR, 926-68071, 1:10000).

#### Transmission Electron Microscopy

Cells were fixed with 0.1 M sodium cacodylate buffer (pH 7.4) containing 2.5% glutaraldehyde, 1.25% paraformaldehyde, and 0.03% picric acid at room temperature over night. After being washed in 0.1M sodium cacodylate buffer (pH 7.4), the cells were then postfixed for 30 min in 1% osmium tetroxide (OsO4)/1.5% potassium ferrocyanide (KFeCN6). These cell samples were washed in water for 3 times and incubated in 1% aqueous uranyl acetate for 30 min followed by another 2 washes in water and subsequent dehydration in different grades of alcohol (5 min each; 50%, 70%, 95%, twice of 100%). Cells were removed from the dish in propylene oxide, pelleted at 3000 rpm for 3 min, and infiltrated for 2 hours to over night in a mixture of propylene oxide and TAAB Epon (1:1 ratio) (Marivac Canada Inc.). The samples were subsequently embedded in TAAB Epon and incubated at 60 °C for polymerization (48 hours). Ultrathin sections were cut on a Reichert Ultracut-S microtome, transferred to copper grids, and stained with lead citrate. Representative micrographs were recorded with an AMT 2k CCD camera in a JEOL 1200EX transmission electron microscope.

#### Confocal Microscopy

Live cell imaging of U2OS expressing GFP-calreticulin was performed on glass-bottom dishes (MatTek) using an Olympus FV1200 laser scanning confocal microscope (Olympus America Inc.). GFP was excited by 473 nm diode laser and emission was captured with band pass filters of 490-540 nm. All representative images were processed following community standards and correctly represented the original data using ImageJ.

#### Secreted Protein Analysis

Three million WM793 cells were seeded in 20 ml RPMI to each 15 cm dish on day 0 (2-4 plates for each condition, 0 h). Since the following treatments initiated ER dilation with different kinetics, they were started at different time points but were synchronized at the point of media replacement on day 3 so that similar levels of ER dilation (60%-90%) were achieved during collection of secreted proteins (8 h of collection time). The treatments included brefeldin A (1 μM, 28-36 h, started on day 1), 6AN (20 μM, 40-48 h, started on day 1), and NMS873 (5 μM, 14-22 h, started on day 2). The old media was carefully aspirated from plates and then warm PBS was used to wash cells twice. With 10 ml fresh warm RPMI (without FBS but still containing compounds) added per plate, the cell secretions were collected 8 hours later. After centrifuging at 1000g for 10 min to pellet the debris, the supernatant was concentrated with Amicon spin filters (10kDa cutoff, MilliporeSigma) according to manufacturer’s instructions. BCA assays were used to quantify the protein concentrations before loading the same amount of protein for SDS-PAGE. Routine in-gel digestion, TMT labeling, and LC-MS analysis was performed.

#### Proteomic Sample Preparation and LC-MS

The cell pellets were lysed in 150 µl of protein extraction buffer superB (Covaris) and were processed on the Covaris S220 shearing device for 180 seconds. The lysates were transferred to empty tubes for reduction, alkylation, and ice-cold chloroform/methanol precipitation. The protein precipitates were transferred to clean Eppendorf tubes with accurately measured weights. After drying, the tubes were weighed again to obtain sample quantities. 200 µg of each protein sample was used for FASP type of digestion and TMT™ (Thermo) labeling according to manufacturer’s instructions.

The labeled peptides were separated on Agilent (Santa Clara, CA) 1100 HPLC system using PolyWAX LP column (200×2.1 mm, 5 μm, 300 Å, PolyLC, Columbia MD) for Electrostatic Repulsion Hydrophilic Interaction Chromatography (ERLIC). Peptides were separated across a 90 min gradient from 100% buffer A (90% acetonitrile, 0.1% acetic acid) to 75 % buffer B (30% acetonitrile, 0.1% formic acid) with 20 fractions collected by time. Each fraction was dried in SpeedVac (Eppendorf, Germany) and re-suspended in 0.1% formic acid solution before injection to a mass spectrometer.

Each fraction was submitted for single LC-MS/MS experiment that was performed on a Tribrid Lumos Orbitrap (Thermo) equipped with an EASY1000 HPLC nano pump. Peptides were separated using a microcapillary trapping column (100 µm inner diameter) in-house packed with approximately 5 cm of C18 Reprosil resin (5 µm, 100 Å, Dr. Maisch GmbH, Germany) followed by analytical column ~25 cm of Reprosil resin (1.9 µm, 200 Å, Dr. Maisch GmbH, Germany). Separation was achieved through applying a gradient from 5–27% ACN in 0.1% formic acid over 90 min at 150 nl min^−1^. Electrospray ionization was enabled through applying a voltage of 2 kV using a home-made electrode junction at the end of the microcapillary column and sprayed from fused silica pico tips (New Objective, MA). The Tribrid Lumos Orbitrap was operated in data-dependent mode. The mass spectrometry survey scan was performed in the Orbitrap with a range of 400– 1,800 m/z at a resolution of 6 × 10^4^. The top 20 ions were subjected to HCD MS2 event in Orbitrap part of the instrument. The fragment ion isolation width was set to 0.7 m/z, AGC was set to 50,000 ions, the maximum ion time was 150 ms, normalized collision energy was set to 37V and an activation time of 1 ms for each HCD MS2 scan with resolution of 5×10^4^, same ions were submitted in parallel into ion trap part of the instrument for CID type of fragmentation to improve identification rate.

#### Proteomic Data Analysis and Quantification

Raw data were submitted for analysis in Proteome Discoverer 2.1.0366 (Thermo Scientific) software. Assignment of MS/MS spectra was performed using the Sequest HT algorithm by searching the data against a protein sequence database including all entries from the Human Uniprot database (SwissProt 16,792, 2016) and other known contaminants including human keratins and common lab contaminants. Sequest HT searches were performed using a 15 ppm precursor ion tolerance and requiring each peptide terminus to follow trypsin protease specificity, while allowing up to two missed cleavages. 10-plex TMT tags on peptide N-termini and lysine residues (+229.162932 Da) were set as static modifications while methionine oxidation (+15.99492 Da) was set as a variable modification. A MS2 spectra assignment false discovery rate (FDR) of 1% on protein level was achieved by applying the target-decoy database search. Filtering was performed using a Percolator (Käll et al., 2008). For quantification, a 0.02 m/z window centered on the theoretical m/z value of the reporter ions was applied and the intensity of the signal closest to the theoretical m/z value was recorded. Reporter ion intensities were exported in result file of Proteome Discoverer 2.1 search engine as excel tables. The total signal intensity across all peptides quantified was summed for each TMT channel, and all intensity values were adjusted to account for potential sample handling variance in each labeled channel.

#### Gene Set Enrichment Analysis (GSEA)

For the whole proteome TMT quantitation data from WM793, two sample t-tests were performed to compare the treated groups versus control groups. Proteins with possible up-regulations or down-regulations (filtered with p<0.05) in each dataset were pre-ranked by their fold changes. C5 gene ontology cellular component sets (c5.cc.v6.2) from the Molecular Signatures Database (http://software.broadinstitute.org/gsea/msigdb) were chosen for this analysis (minimum size: 10, maximum size: 500). The *fgsea* R package (Sergushichev, 2016) was then applied to calculate normalized enrichment scores (NES) and other metrics based on the selected proteins.

### QUANTIFICATION AND STATISTICAL ANALYSIS

All statistical analyses used in this paper were done in R v 3.4.2 (downloaded from https://www.r-project.org/). Data visualization was done in R and Prism (GraphPad). Statistical tests (two-sided) including the number of sample biological replicates (n) and statistical significance (p) are reported in the figures and associated legends. If applicable, we used the Benjamini-Hochberg procedure to control for multiple hypothesis testing.

### DATA AND SOFTWARE AVAILABILITY

The data and related analyses can be found in Tables S1-2. Additional CCLE resources are available at: https://portals.broadinstitute.org/ccle.

---

**Figure S1.**
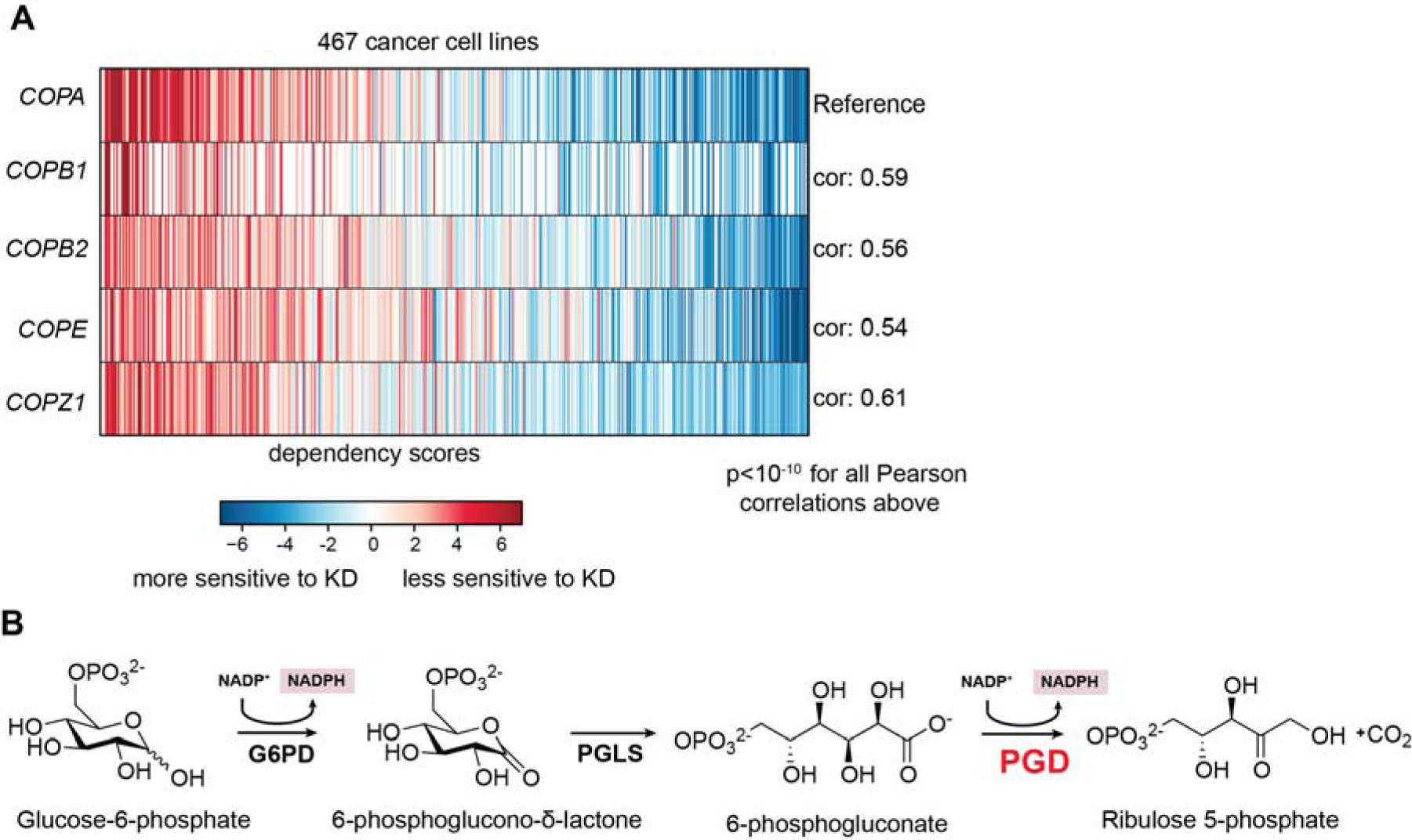
PGD as a Metabolic Dependency, Related to Figure 1. (A) Heatmap representation of the coatomer complex dependency revealed in the RNAi-screen including 467 cancer cell lines. *COPA*, *COPB1*, *COPB2*, *COPE*, and *COPZ* are available in the dataset. DEMETER scores are used as quantitative metrics to show cell line dependencies on specific genes (related to viabilities after gene knockdown). (B) Schematic showing the oxidative phase in pentose phosphate pathway.

**Figure S2.**
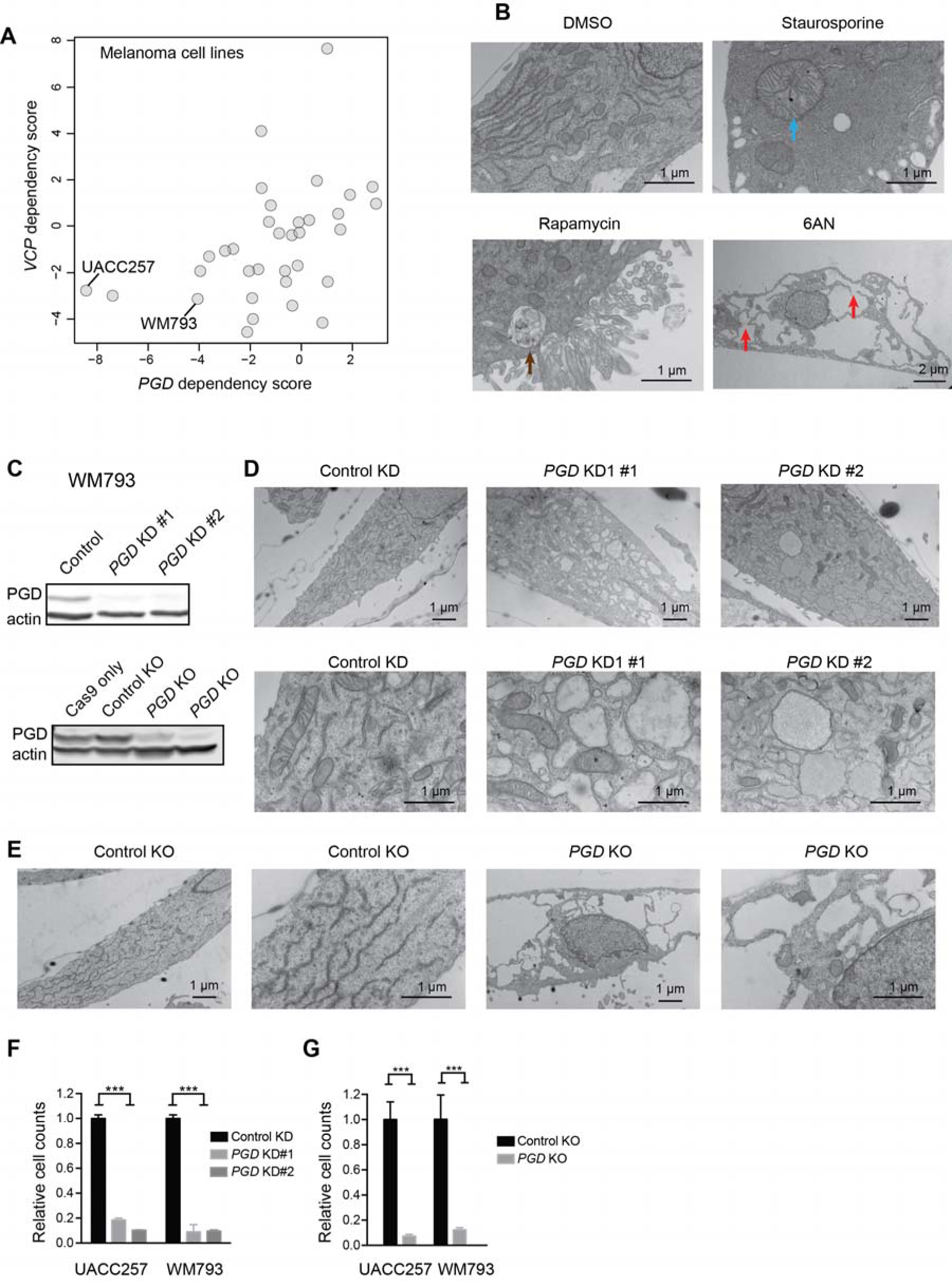
PGD Suppression Induces a Distinct Stress Phenotype Characterized by ER Dilation, Related to Figure 2. (A) Comparisons of *PGD* dependency and *VCP* dependency in melanoma cell lines. The model cell lines are labeled. (B) Representative TEM micrographs comparing different cell death or stress phenotypes using UACC257 cells. An abnormal mitochondrion is noted with a blue arrow. An autophagic vacuole is noted with a brown arrow. Sites where ER vacuoles might be merging are noted with red arrows. The treatments include staurosporine (1 μM, 24h), rapamycin (100 nM, 24h), and 6AN (20 μM, 72h). (C) Top, an immunoblot of PGD in WM793 cells with *PGD* KD or control KD. Bottom, an immunoblot of PGD in WM793 cells with doxycycline-induced *PGD* KO or control KO. Actin was used as the loading control in both. (D) Representative TEM micrographs showing impact of *PGD* KD using WM793 cells as an example. Top, lower magnification; bottom, higher magnification. (D) Representative TEM micrographs showing impact of *PGD* KO using WM793 cells as an example. (F-G) Impact of PGD KD (F) or KO (G) on cell proliferation. The cells were counted 1 week after PGD KD or induced KO. Bar plots represent mean ± SEM (n=3) (*** p < 0.005).

**Figure S3.**
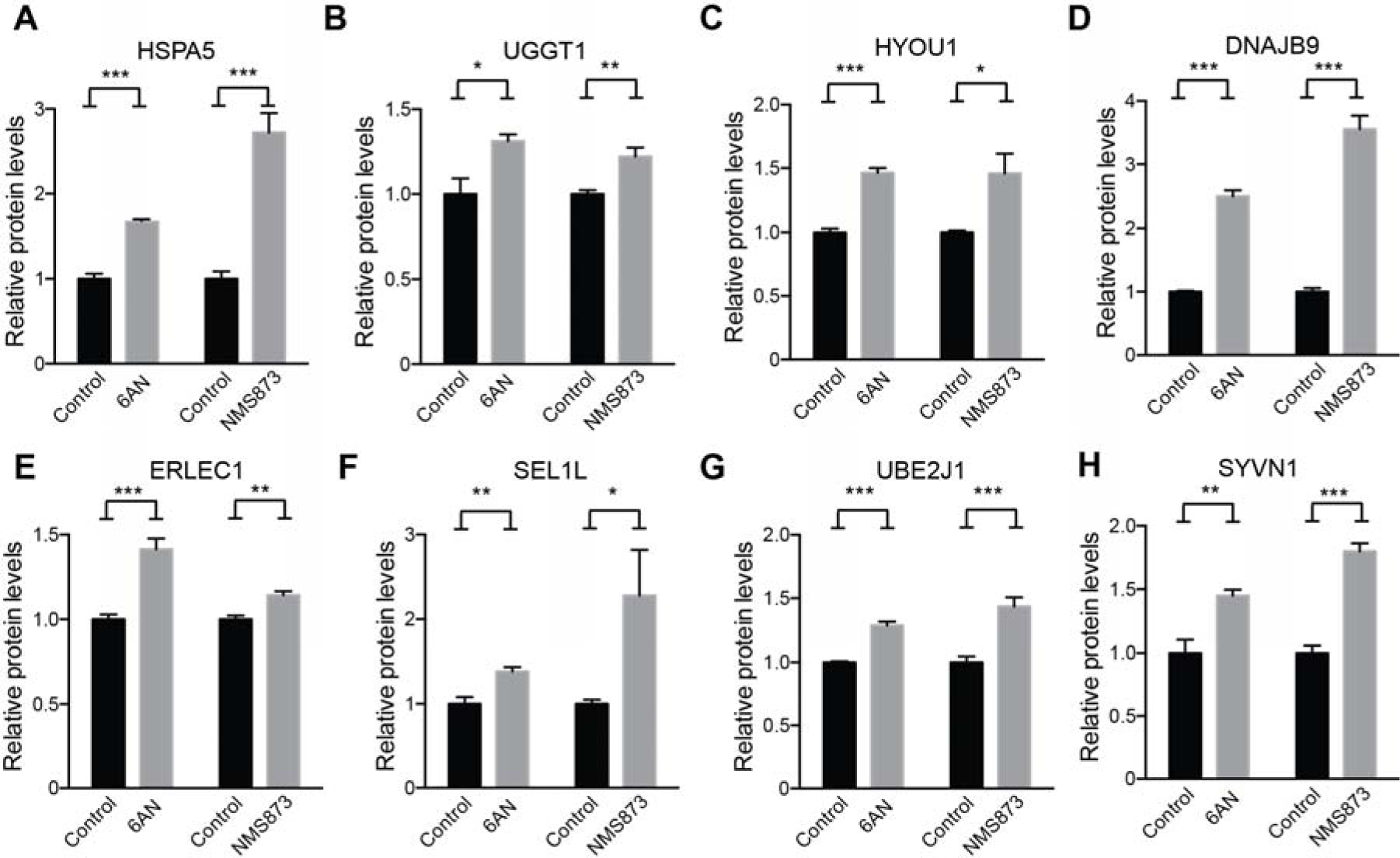
Key Machineries Involved in Misfolded Protein Folding and Export are Up-regulated in Cells with ER Dilation, Related to Figure 3. (A) Relative protein levels of (A) HSPA5, (B) UGGT1, (C) HYOU1, (D) DNAJB9, (E) ERLEC1, (F) SEL1L, (G) UBE2J1, (H) SYVN1 based on quantitative proteomic measurements. The bar plots represent mean ± SEM (*** p < 0.005; ** p < 0.01; * p < 0.05; n = 3).

**Figure S4.**
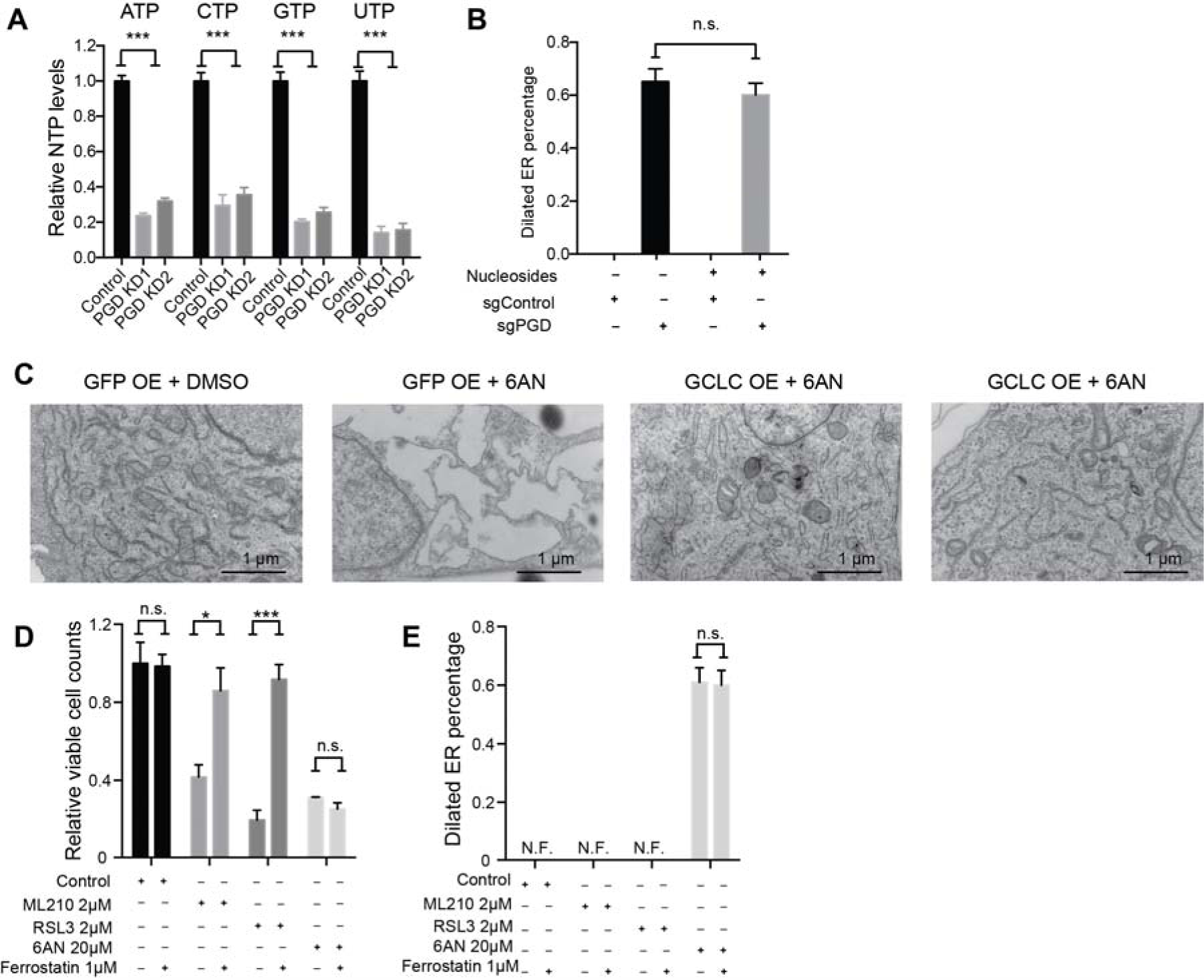
Rescues of ER Dilations Induced by PGD Suppression, Related to Figure 4. (A) Relative nucleoside triphosphate levels in WM793 cells with *PGD* KD versus control. (B) Effects of nucleoside supplements (30 μM cytidine, guanosine, uridine, adenosine, and thymidine) on *PGD* KO-induced ER dilation in WM793 cells (9 days after 1 μg/ml doxycycline treatment). (C) Representative TEM micrographs showing impact of *GCLC* overexpression in WM793 cells when treated with 6AN. (D) Effects of ferrostatin on viable cell counts 2 days after treatment of ferroptosis inducers or 6AN in WM793. (E) Same as (D) but estimating percentages of ER dilation. The bar plots represent mean ± SEM (*** p < 0.005; ** p < 0.01; * p < 0.05; n.s., non-significant, p > 0.05; n = 4) (A, B, D, E).

